# Conformational and Functional Regulation of SET by Legumain Cleavage

**DOI:** 10.1101/2025.01.28.635311

**Authors:** Carina Horak, Alexander C. Wieland, Rupert Klaushofer, Peter Briza, Hans Brandstetter, Elfriede Dall

## Abstract

The cysteine protease legumain typically localizes to the endolysosomal system, where it is an important player in the immune system. However, in the context of Alzheimer’s disease (AD), legumain has been shown to be translocated to the cytosol, where it cleaves SET, synonymously termed TAF-1 or I2PP2A, an inhibitor of protein phosphatase 2A. SET is primarily found in the nucleus, where it regulates gene transcription, cell cycle progression, and histone acetylation, but can also translocate to the cytoplasm where it regulates cell migration and is implicated in neuronal apoptosis in AD. In this study, we demonstrate that legumain cleaves SET at two major sites: Asn16 at the N-terminal end and Asn175 at the earmuff domain. Contrary to previous findings, our biochemical and crystallographic experiments reveal that the corresponding N- and C-terminal cleavage products remain bound in a stable complex, rather than dissociating. Additionally, we show that the C-terminal acidic stretch of SET is essential for its binding to histone 1, and that cleavage impairs this interaction. Finally, we demonstrate that SET positively modulates PP2A activity. This effect is however abolished upon cleavage by legumain.

## Introduction

Proteases are important signaling molecules that are involved in virtually all biological processes in all living organisms. By consequence, dysregulation of their protease activity can have severe effects resulting in a variety of pathologies, including cancer and Alzheimer’s disease (1,2). Therefore, proteases must be regulated delicately and on different levels. The cysteine protease legumain is an important player of our immune system.

Although the human cysteine protease legumain is structurally homologous to the cytosolic caspases, the key mediators of apoptosis, human legumain is localized primarily to the acidic endo-lysosome (3). Legumain’s best established function is the processing of self (myelin basic protein; MBP), limiting tolerance to the autoantigen, and foreign (tetanus toxin C-terminal fragment; TTCF) proteins, promoting presentation on the MHCII complex. Moreover, it is proteolytically activating Toll-like receptors, and endo-lysosomal cysteine cathepsins (4–6). Because of its strict specificity for cleaving after asparagine residues, it is synonymously termed the asparaginyl endopeptidase (AEP) (7,8). On a pathophysiological level, legumain is overexpressed in the majority of human solid tumors, including breast cancer, and colorectal cancer (9,10). Here, legumain is associated with enhanced tissue invasion and metastasis and as a consequence overexpression correlates with poor prognosis. Under these conditions legumain was found translocated to the nucleus, cytoplasm and the extracellular space (9,11–14). Recently, there is growing evidence that legumain is similarly translocated in the aged brain and thereby facilitates aggregation of proteins critically linked with neurodegenerative diseases, finally causing neuronal damage (15–18). In the context of Alzheimer’s disease (AD), cytosolic legumain has been shown to process I_2_^PP2A^, an inhibitor of protein phosphatase 2A (PP2A), which plays a critical role in AD (19). I_2_^PP2A^, also known as SET (which we will use throughout this manuscript) or TAF-1, is a 39-kDa protein predominantly localized in the nucleus but capable of translocating to the cytoplasm, where it induces cell migration (20–22). SET also regulates gene transcription (23), cell cycle progression (24), and histone acetylation (25), and is implicated in neuronal apoptosis in AD (26). Notably, in AD brains, SET is selectively cleaved at Asn175 (27). Gustavo Basurto-Islas and colleagues demonstrated that cytosolic legumain mediates this cleavage, resulting in the activation of SET. Upon cleavage at Asn175, SET inhibits PP2A activity, leading to tau hyperphosphorylation and subsequent formation of neurofibrillary tangles (NFTs), composed of hyperphosphorylated tau fragments (11).

Despite these findings, the molecular details of SET cleavage and its structural consequences remain poorly understood. Previous studies suggest that legumain-mediated cleavage results in dissociation into separate N- and C-terminal SET cleavage products, both of which may exhibit inhibitory functions. However, this hypothesis has not been directly tested with isolated proteins. Therefore, within this study we set out to investigate the structural and functional effects of SET cleavage by legumain.

## Results

### Legumain cleaves SET at two major sites

We successfully established the recombinant expression of full-length SET (SET-FL, 2–277) and a C-terminally truncated version, which lacks the predicted unstructured acidic C-terminal region (SETΔC, 2–225), in *E. coli* (Fig. 1). Subsequently, cleavage by legumain was tested at pH 5.5, which corresponds to the pH optimum for legumain activity. At a molar ratio of 1:10 (AEP:SET), SET-FL was completely converted into two predominant cleavage products (Fig. 2A). Similarly, truncated SETΔC was also efficiently converted into two cleavage products. Interestingly, the higher molecular weight cleavage product migrated at the same position in both samples, suggesting it represents the N-terminal cleavage product, since the N-terminal region was identical in both constructs. SET-FL and SETΔC differed in their C-terminal region only. Subsequently, we also tested cleavage under near-neutral pH conditions to mimic cytosolic environments. Under these conditions, cleavage also occurred, but with lower efficiency compared to pH 5.5, which is in line with legumain’s reduced stability at near-neutral pH (Fig. 2B). Nevertheless, cleavage at near-neutral pH was still possible.

**Figure 1.**
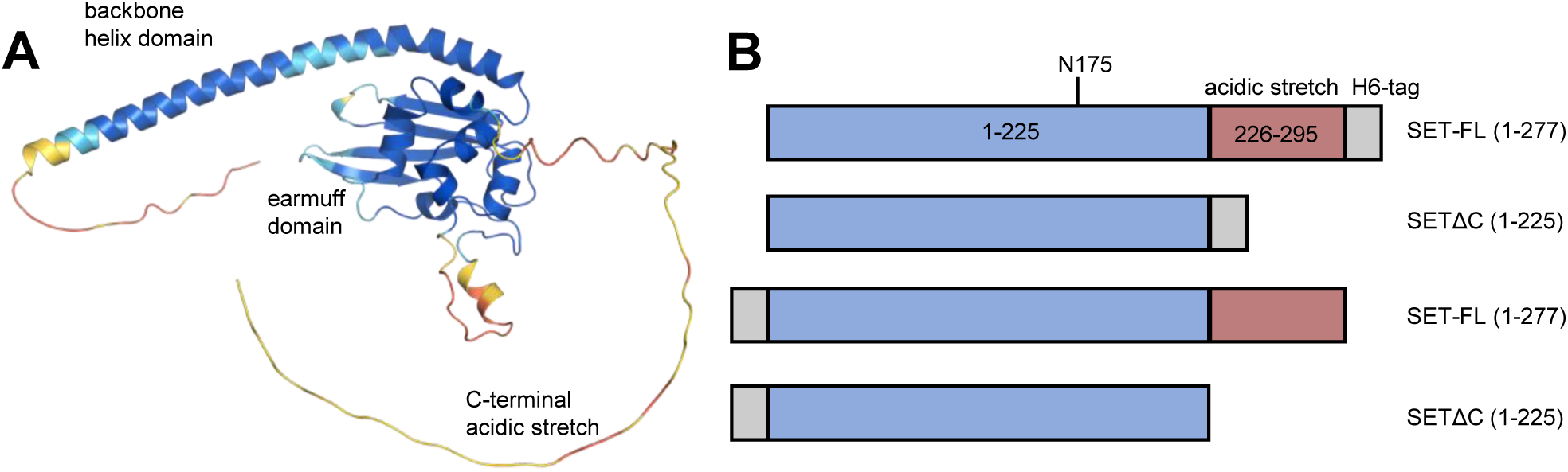
SET is built up by a backbone helix domain, an earmuff domain and a C-terminal acidic stretch. **A)** AlphaFold model of SET-FL, color-coded according to the per-residue predicted local distance difference test (pLDDT) value (blue: high pLDDT, red: low pLDDT). The AlphaFold run resulted in 5 separate models. The model with the highest overall pLDDT (pLDDT: 78.9) is shown. **B)** Scheme of constructs used in this study.

**Figure 2.**
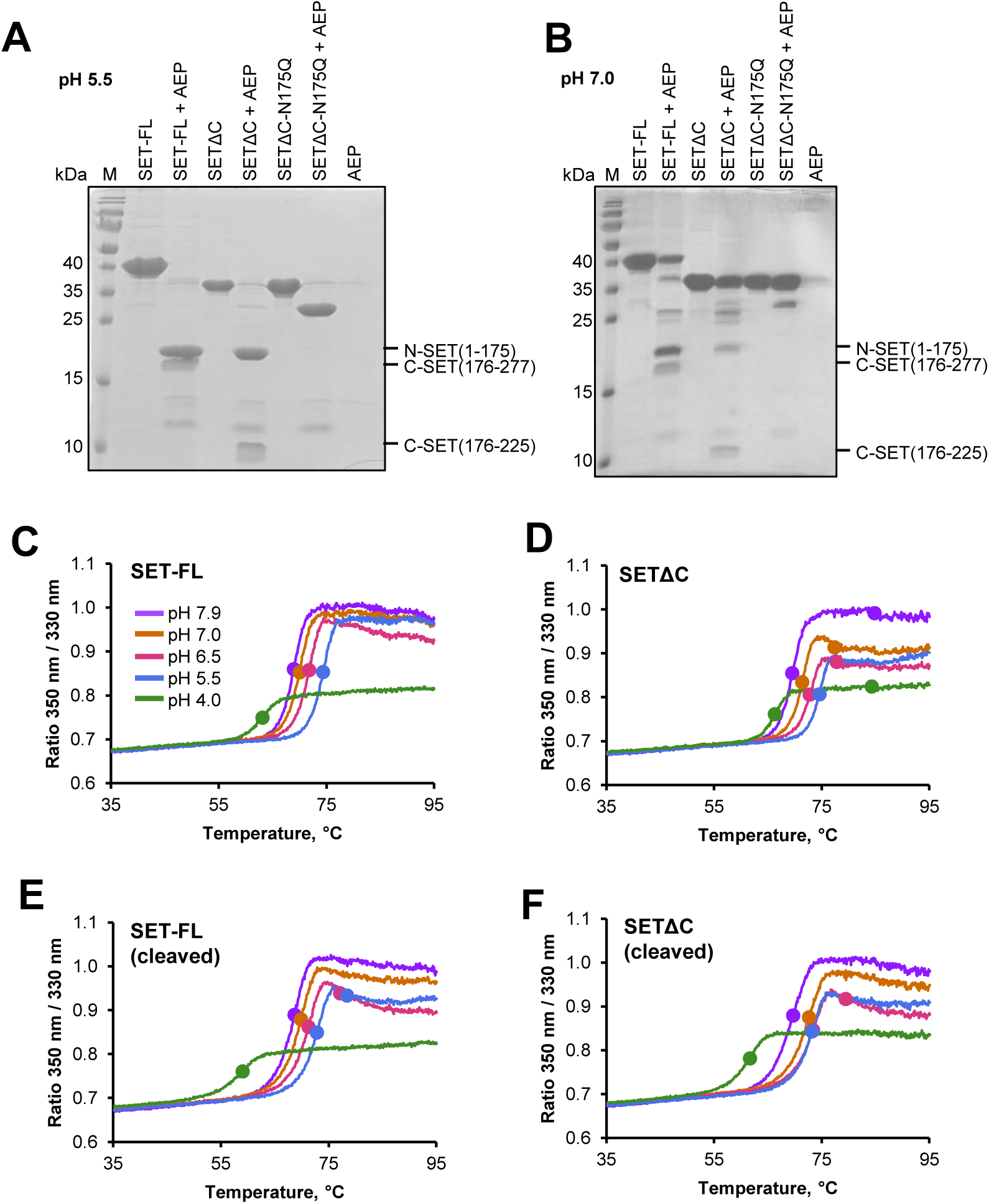
Cleavage of SET by legumain is pH-dependent. **A)** and **B)** SET-FL, SETΔC and SETΔC-N175Q constructs were incubated with legumain at indicated pH values for 1.5 hours at 37 °C. **C)**, **D)**, **E)** and **F)** Thermal stability of SET-constructs at indicated pH values measured by nanoDSF experiments. Inflection temperatures are indicated by filled circles.

To rule out the possibility that the pH-dependent processing was due to reduced conformational stability of SET at acidic pH, we examined its pH-dependent thermal stability using nanoDSF experiments (Fig. 2C-F). Interestingly, SET demonstrated a high thermal stability (>65 °C) across all tested pH conditions, with the highest stability observed at pH 5.5 (Table 1). Therefore, the observed pH-dependent difference in processing could not be attributed to changes in conformational stability, but rather is likely due to the lower pH stability of legumain at near-neutral pH. Importantly, SET-FL and SETΔC showed similar inflection temperatures before and after cleavage (Fig. 2C-F and Table 1). Highlighting that cleaved SET was still conformationally stable.

**Table 1.** SET is most stable at pH 5.5. Inflection temperatures (T_i_) of SET constructs at indicated pH values determined in nanoDSF experiments.

| pH | $T_i$ SET-FL (°C) | $T_i$ SET-FL cleaved (°C) | $T_i$ SETΔC (°C) | $T_i$ SETΔC cleaved (°C) |
| --- | --- | --- | --- | --- |
| 8.0 | 69.0 | 68.7 | 69.5 | 69.7 |
| 7.0 | 70.0 | 69.9 | 71.4 | 72.7 |
| 6.5 | 71.8 | 71.3 | 72.8 | 73.4 |
| 5.5 | 74.4 | 72.8 | 74.6 | 73.2 |
| 4.0 | 63.1 | 59 | 66.3 | 61.7 |

To identify the cleavage sites, we performed mass spectrometry analysis, which revealed two major sites: Asn175 on the earmuff domain, which was also identified in a previous study (11), and Asn16 on the N-terminal end of the SET protein (Suppl. Fig. S1 and S2). To confirm these sites, we prepared an N175Q-SET point mutant and tested its cleavage by legumain. As expected, this mutant only showed cleavage at the N-terminal end of the protein (Fig. 2A,B and Suppl. Fig. S3).

### SET remains intact after cleavage

Next, we aimed to investigate how cleavage of SET affects its molecular structure and biophysical properties. One key question was whether SET dissociated into separate N- and C-terminal cleavage products following cleavage, as previously suggested (11). To explore this in more detail, we performed size-exclusion chromatography (SEC) experiments using uncleaved and cleaved SET proteins. Importantly, we observed no shift in the elution volume when comparing uncleaved and cleaved SET proteins (Fig. 3 and Suppl. Fig. S4). Additionally, SDS-PAGE analysis of the SEC fractions showed that the cleavage products co-eluted within the same SEC peak. This finding was consistent for both full-length SET and the C-terminally truncated SET (SETΔC) constructs. Suggesting that the cleavage products did not dissociate into separate N- and C-terminal fragments.

**Figure 3.**
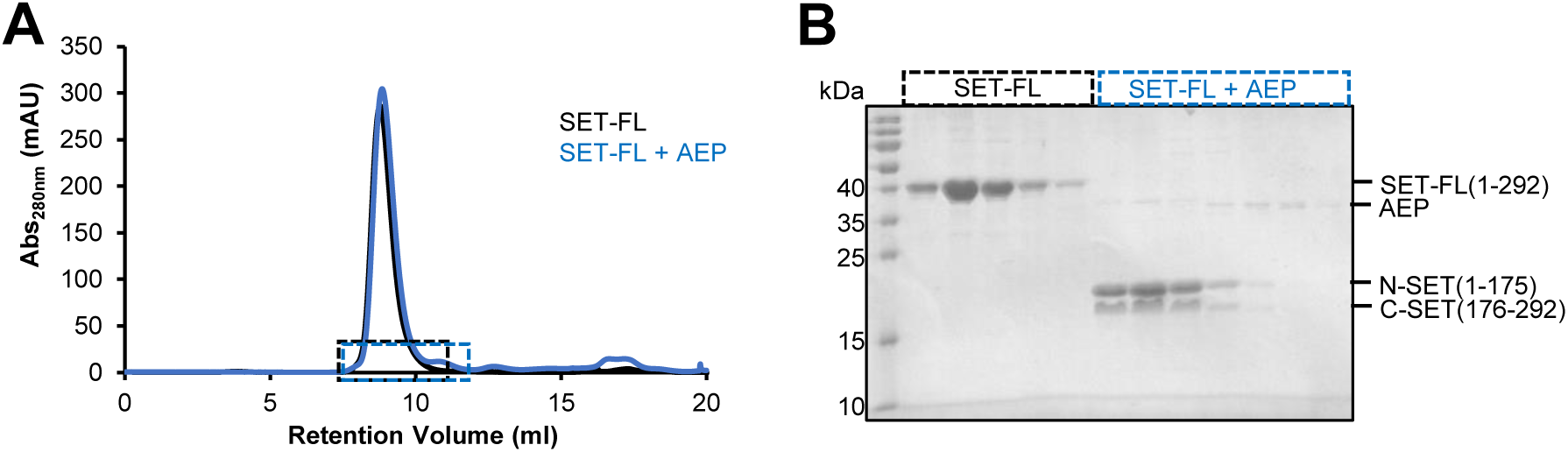
Cleavage products of SET-FL do not dissociate after cleavage. **A)** Size exclusion chromatography experiment of intact and cleaved SET-FL at pH 5.5. **B)** Peak fractions of the experiment shown in E) were analyzed by SDS-PAGE.

To further confirm whether SET really remains intact after cleavage and to examine any potential conformational changes caused by cleavage, we crystallized and solved the crystal structure of cleaved SETΔC (Table 2). Remarkably, the full-length protein was observed in the crystal structure (Fig. 4). SETΔC did not dissociate into separate N- and C-terminal fragments. Cleaved SETΔC was similarly built up by a backbone helix domain and an earmuff domain (Fig. 4A). Importantly, the short N-terminal helix present in uncleaved SETΔC (pdb 2e50) (28), was absent in cleaved SETΔC because it was processed after Asn16. Asn16 was located on the linker between the N-terminal helix and the backbone helix domain. When comparing the structures of uncleaved and cleaved SETΔC, we found them to be highly similar, with an overall RMSD of 0.8 Å for the Cα atoms. However, while the earmuff domains showed an RMSD of only 0.45 Å, the backbone helix domains superimposed with a higher RMSD of 1.19 Å. Both structures showed monomers assembled into a C2-symmetric dimer with a “headphone-like” shape (Fig. 4B). Importantly, the assembly of the monomers in the dimer differed significantly between the two, with an RMSD of 4.0 Å (Fig. 4C). The Asn175 cleavage site was located on a flexible loop on the earmuff domain which was therefore not visible in the electron density map (Fig. 4D). Importantly, this loop was more rigid in the structure of cleaved SETΔC compared to uncleaved SETΔC. Altogether, these finding suggested that cleavage induced subtle conformational changes that may affect the protein’s function.

**Figure 4.**
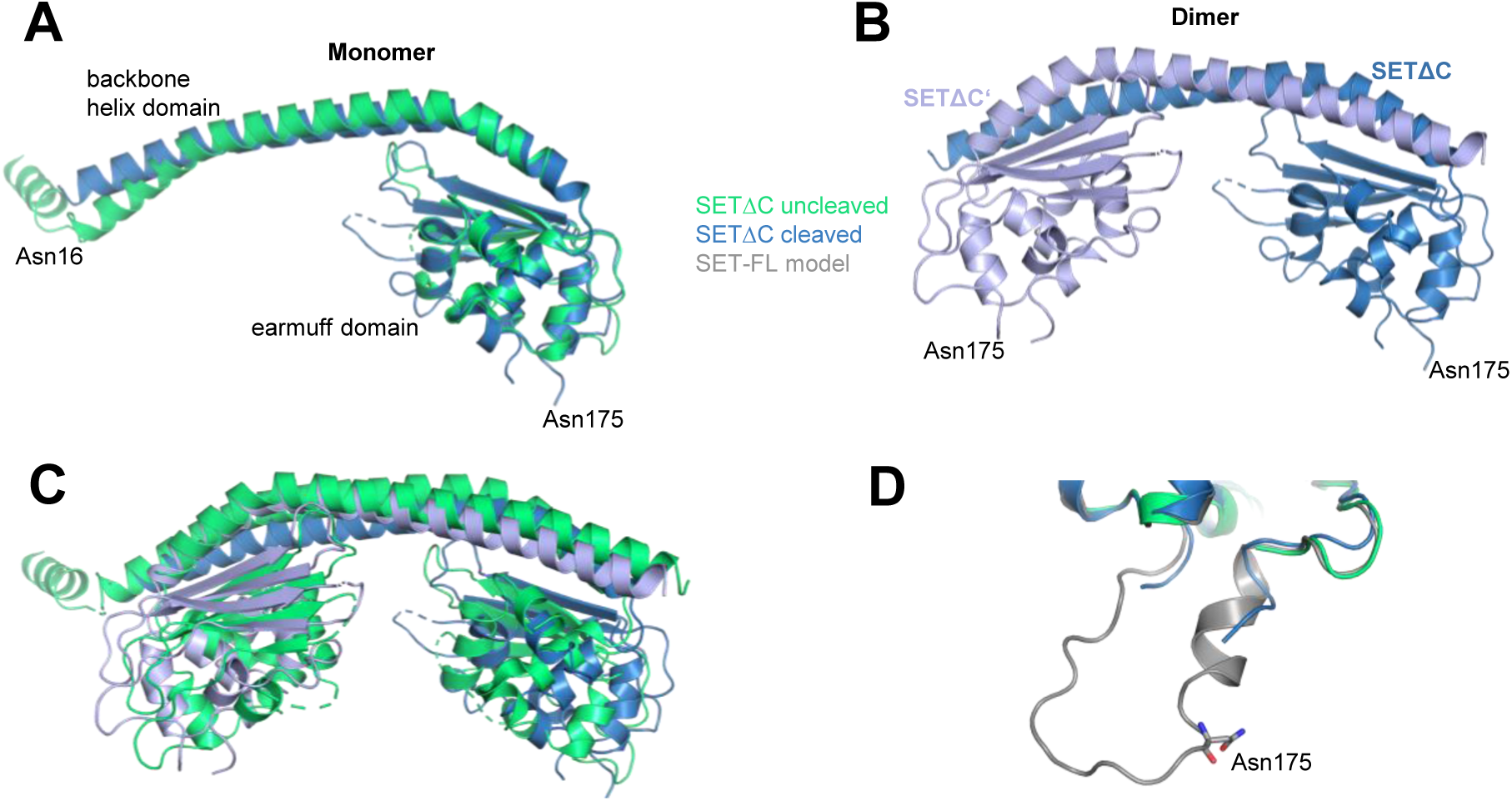
Crystal structure of SETΔC after cleavage by legumain. **A)** Superposition of cleaved (blue) and intact (green, pdb 2e50) SETΔC monomers. **B)** Cleaved SETΔC showed a dimeric assembly in the crystal structure. The Asn175 cleavage site was located on a flexible loop which was not resolved in the electron density map. **C)** Superposition of cleaved and intact SETΔC dimers. **D)** Zoom-in on the Asn175 legumain cleavage site. Asn175 is shown as sticks in the AlphaFold model of SET-FL.

**Table 2.**
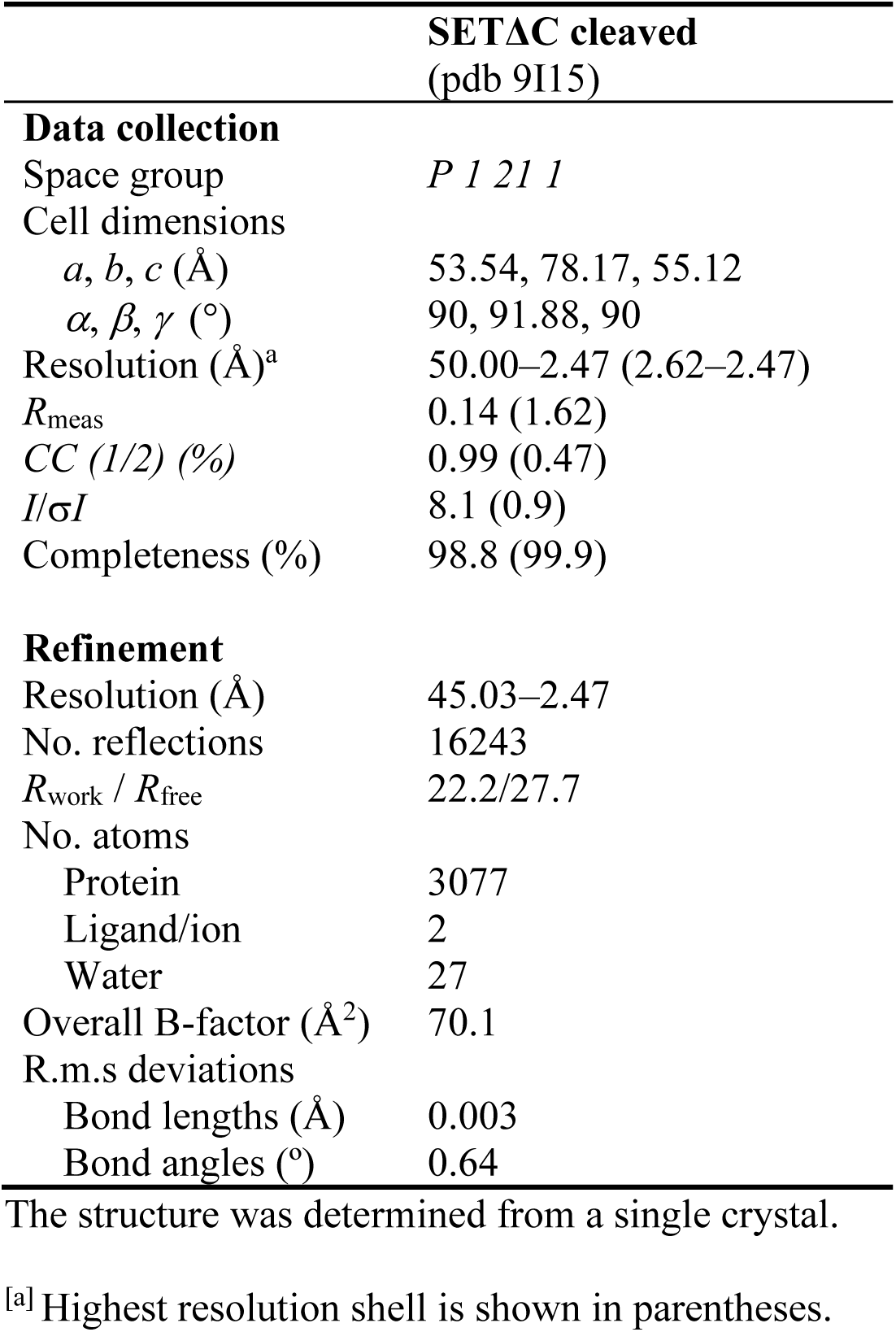
Xray data collection and refinement statistics.

|  | **SETΔC cleaved**<br>(pdb 9I15) |
| --- | --- |
| **Data collection** |  |
| Space group | *P 1 21 1* |
| Cell dimensions |  |
| *a*, *b*, *c* (Å) | 53.54, 78.17, 55.12 |
| $\alpha$, $\beta$, $\gamma$ (°) | 90, 91.88, 90 |
| Resolution (Å)<sup>a</sup> | 50.00–2.47 (2.62–2.47) |
| *R*<sub>meas</sub> | 0.14 (1.62) |
| *CC* (1/2) (%) | 0.99 (0.47) |
| *I*/σ*I* | 8.1 (0.9) |
| Completeness (%) | 98.8 (99.9) |
| **Refinement** |  |
| Resolution (Å) | 45.03–2.47 |
| No. reflections | 16243 |
| *R*<sub>work</sub> / *R*<sub>free</sub> | 22.2/27.7 |
| No. atoms |  |
| Protein | 3077 |
| Ligand/ion | 2 |
| Water | 27 |
| Overall B-factor (Å<sup>2</sup>) | 70.1 |
| R.m.s deviations |  |
| Bond lengths (Å) | 0.003 |
| Bond angles (°) | 0.64 |

The structure was determined from a single crystal.
<sup>[a]</sup> Highest resolution shell is shown in parentheses.

### Binding to histone 1 is modulated upon cleavage

To determine whether the structural differences between cleaved and uncleaved SET translated into functional changes, we tested two of SET’s main functions: histone binding and PP2A inhibition. For histone binding, we incubated our recombinant SET constructs with a mix of bovine histones and identified the interacting proteins using pull-down experiments. Specifically, we employed the C-terminal His_6_-tag on our SET constructs for Ni²⁺-affinity purification after incubation with the histone mix. Any histones that interacted with SET should consequently co-elute during this process. In a control experiment, we tested whether the histones alone could bind to the Ni²⁺-resin. We found that histones did not bind to the Ni²⁺-resin in the absence of SET, while SET-FL and SETΔC did bind (Suppl. Fig. S5). When we co-incubated SET-FL with the histone mix, all histones co-eluted with SET-FL, indicating that all histones interacted with the full-length protein (Fig. 5A, left). However, when we co-incubated histones with SETΔC, all core histones (histones H2A&B, H3 and H4) interacted with the truncated protein but the linker histone H1 did not (Fig. 5B). This suggested that histone H1 binds specifically to the C-terminal acidic region of SET. In subsequent experiments with cleaved SET variants, we observed a reduction in the interaction between histone H1 and SET-FL, indicating that cleavage affects the binding of histone H1 (Fig. 5A, right).

**Figure 5.**
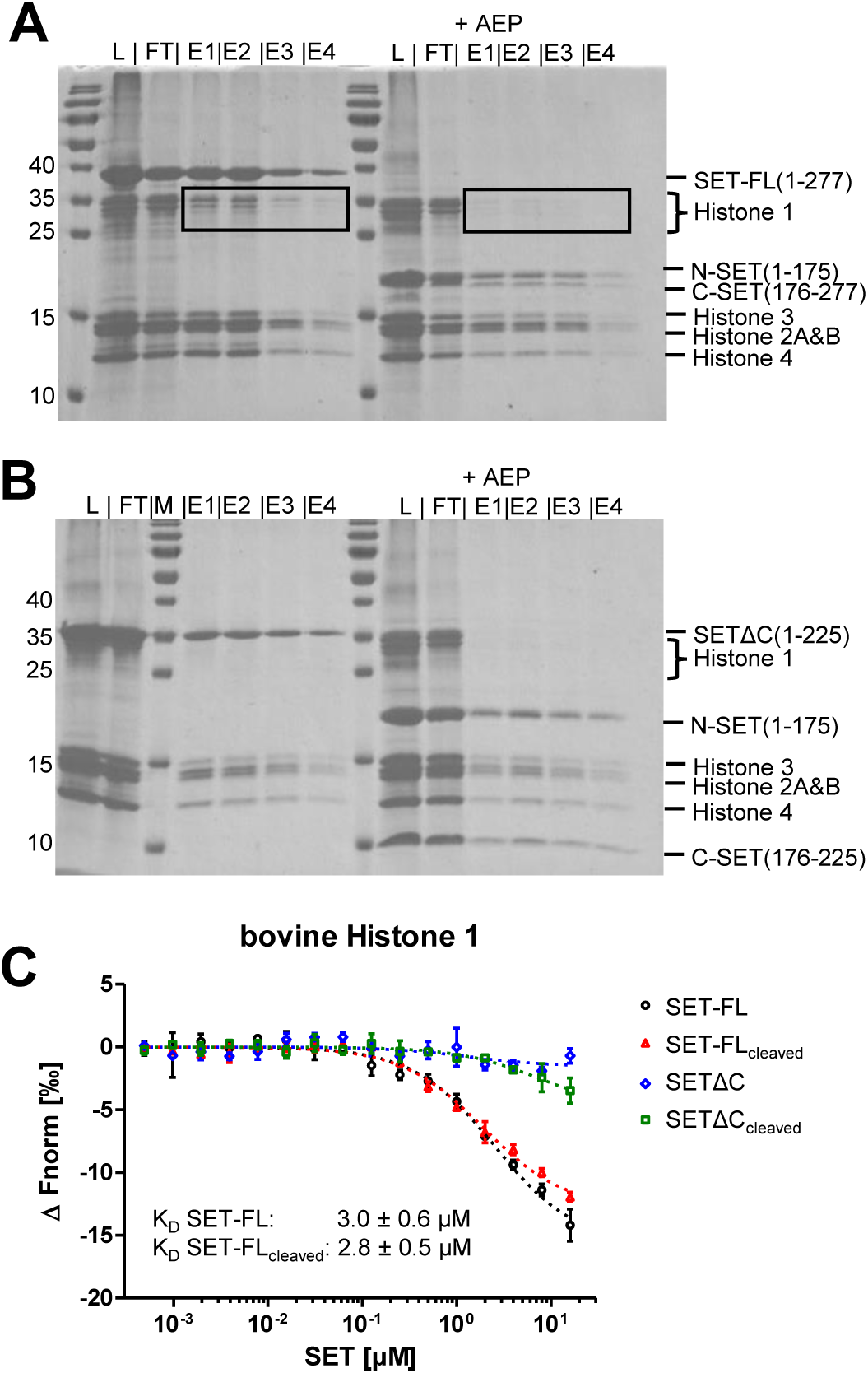
Uncleaved and cleaved SET bind to histones. **A)** and **B)** Pull-down experiments of SET-FL and SETΔC after incubation with a mix of bovine histones. Histones that bound to SET were co-eluted after Ni^2+^-purification, exploiting the C-terminal His_6_-tag of the SET constructs. L: load, FT: flow through, M: molecular weight standard, E1-4: elution fractions 1-4 **C)** Microscale thermophoresis (MST) experiments testing the interaction of indicated SET variants to histone 1.

**Figure 6.**
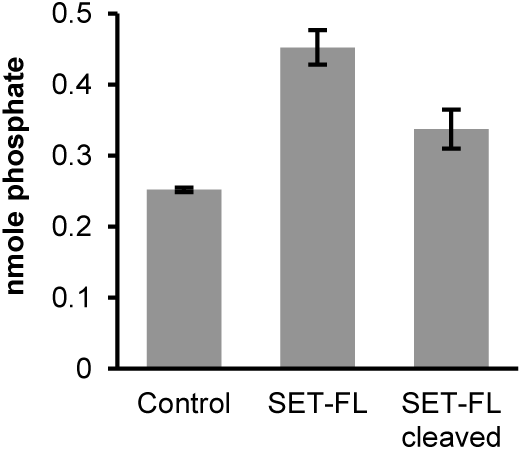
The stimulating effect of SET-FL on PP2A activity is reduced upon cleavage by legumain. Activity of PP2A was measured using a malachite green phosphate assay where the phosphorylated K-R-pT-I-R-R substrate of PP2A is detected as an increase in absorbance at 620 nm.

To further investigate the relevance of the C-terminal acidic region, we purchased bovine histone H1 and tested its interaction with both uncleaved and cleaved SET-FL using Microscale Thermophoresis (MST) experiments. Interestingly, we did not observe a significant difference in binding affinity between uncleaved SET-FL and cleaved SET-FL (Fig. 5C). Both showed a K_D_ value of approx 3 µM. Similarly, we investigated the binding of uncleaved and cleaved SETΔC to bovine histone H1. In line with our pull-down experiments, we did not observe significant binding to bovine histone H1 (Fig. 5C). Next, we also analyzed the binding of human histone H1 to our human SET-FL constructs. Interestingly we did observe binding in the micromolar range, however the MST signal was very low and, for this technical reason, could not be used to calculate reliable K_D_ values (Suppl. Fig. S6). Taken together, these results suggest that histone H1 likely forms a complex with other histones, and that it is the binding of this complex that is influenced upon cleavage of SET-FL by legumain.

### Intact SET has an agonistic effect on PP2A activity, which is abolished upon cleavage

Next, we investigated whether cleavage of SET affects its ability to inhibit PP2A activity. To this end, we established a phosphatase assay in which we isolated PP2A from HEK293T cell lysates using immobilized antibody affinity chromatography. We then added a PP2A substrate to the reaction and measured phosphate release using malachite green detection solution. Our results confirmed the presence of PP2A activity in the HEK cell lysates (Fig. 5). Interestingly, when we added SET-FL or cleaved SET-FL proteins prior to the addition of the PP2A substrate, we observed an increase in PP2A activity. However, this increase was less pronounced in the presence of cleaved SET-FL, suggesting that SET has a stabilizing effect on PP2A, which is lost upon cleavage. This assay was repeated several times with qualitatively similar results. It has to be mentioned however, that the magnitude of the effect varied between replicates.

## Discussion

The crystal structure of cleaved SETΔC revealed that, while its overall conformation remained superficially similar to uncleaved SETΔC, subtle structural differences suggest a potential regulatory role for cleavage in modulating SET function.

Indeed, in our histone pull-down experiments, we observed weaker binding of histone 1 to SET-FL following legumain processing. While both SET-FL and SETΔC bound to the core histones (H2A, H2B, H3, and H4), only SET-FL bound convincingly to the linker histone H1 (Fig 5). These findings contradict the results of Muto et al. (28), who showed that only SET-FL binds to all four core histones, whereas SETΔC only binds to histone H3 and H4. This discrepancy could be explained by the use of different histones and SET constructs in the respective experiments. Karetsou et al. demonstrated that SET-FL bound histone H2B and H3 of calf thymus. Additionally, they showed that SET-FL is also capable of binding histone H3-H4 tetramers (29). In general, it can be stated that both uncleaved and cleaved SET-FL and SETΔC were capable of binding histones. However, only SET-FL was capable of binding to histone H1. Therefore, our findings highlight the critical role of the C-terminal acidic stretch of SET in binding to histones, particularly histone 1. Notably, histones can form intermolecular complexes with each other. Not all histones investigated in our study may bind directly to SET—some may associate indirectly through the formation of intermolecular complexes, as for example seen by Karetsou et al. for H3-H4 tetramers (29). Furthermore, in our pull-down experiments, we observed weaker binding of histone 1 to full-length SET following legumain processing. However, MST experiments indicated that both processed and unprocessed SET exhibited similar binding affinities to histone 1. This discrepancy hints at the involvement of another component / histone that was present in the pull-down assay but not in the MST experiments, potentially binding to histone 1 and thereby contributing to an indirect regulatory function of histone 1. Importantly, no binding was detected in the absence of the C-terminal acidic stretch, indicating that this region likely mediates direct interaction, possibly by engaging with the C-terminal basic region of histone 1.

Li et al. previously found that SET is a potent inhibitor of PP2A activity (30). However, another study by Arnaud L. and colleagues suggested that SET only becomes active as PP2A inhibitor after cleavage at Asn175 and subsequent separation into N- and C-terminal SET fragments (31). In contrast, our study shows that SET does not separate into distinct N- and C-terminal fragments but stays non-covalently bound after cleavage by legumain. Furthermore, our PP2A activity assay suggests that SET acts as a stimulator of PP2A activity. This stimulatory effect was however abolished upon cleavage by legumain, suggesting that SET functions as an indirect inhibitor of PP2A activity. One possible mechanism to explain the observed agonistic effect would be that SET-FL has a stabilizing effect on PP2A which is reduced upon cleavage by legumain. It was indeed surprising to see that SET-FL functions as an agonist rather than a direct inhibitor in our PP2A activity assay. However, previous PP2A activity assays were carried out in a different manner and are therefore not directly comparable. Assays using PP2A from HEK293 cells used cell extracts to analyze PP2A activity (11,31). Similarly, PP2A extracted from bovine kidney was used to characterize its inhibition by SET (30,32). In such assays, endogenous SET may still be present and similarly function as an agonist / a stabilizer of PP2A. In the presence of cleaved SET, the reduction in PP2A stabilization might be interpreted as inhibition. Interestingly, in previous studies on the interaction of legumain with cystatins we also uncovered agonistic properties encoded in the legumain inhibitory human cystatin C (33).

## Material and Methods

### Construct design

Expression constructs of full-length SET (SET-FL) were designed based on the sequence deposited in Uniprot under the accession code Q01105-2. A full-length expression construct (Met1 – Asp277) containing an N-terminal His6-tag and TEV recognition sequence was purchased from Thermo Fisher Scientific (Waltham, Massachusetts). The expression construct was subcloned into the pET-28b(+) expression vector using NcoI and XhoI restriction enzyme cleavage sites. Additionally, another construct harboring a C-terminal His6-tag was prepared in the pET-28b(+) expression vector using the same restriction enzyme cleavage sites. For that, a PCR reaction was performed using the primers ATGCCCATGGGTGCACCGGCAGCAAAAGTTAGCAAG (forward, NcoI recognition site) and ATGCCTCGAGATCATCTTCGCCTTCGTCCTCTTCG (reverse, XhoI) for SET-FL. For SETΔC we used the same forward primer but a different reverse primer: ATGCCTCGAGCATATCCGGAACCAGATAATACTGC. The PCR product was purified, digested by the respective restriction enzymes, and ligated into the pET-28b vector. Furthermore, C-terminally truncated constructs of SET (SETΔC) ranging from amino acids Met1 – Met225 were prepared using a similar protocol. Furthermore, SETΔC-N175Q point mutants were purchased form Thermo fisher (Waltham, Massachusetts) harboring an N-terminal His_6_-tag and TEV recognition sequence in the pET-28b(+) expression vector and similarly subcloned to obtain a C-terminal His6-tag following the same procedure as described for the wild-type constructs.

### Expression of SET

The final expression vectors were transformed into the *E. coli* BL21(DE3) strain for expression. Expression flasks of 2 L containing 500 mL of LB medium (Carl Roth LB-Medium, Lennox) and kanamycin at 100 μg/ml concentration were inoculated with 10 ml of preculture and incubated at 37 °C and 230 rpm until an OD600 of approximately 0.8 was reached. Expression was induced upon addition of IPTG (Isopropyl β-D-1-thiogalactopyranoside) at a final concentration of 1 mM. After induction, the cultures were kept overnight at 25 °C and 230 rpm. The cells were harvested and either lysed for immediate use, or frozen as pellets at -20 °C.

### Purification of SET

Cell pellets were resuspended in 20 ml of lysis buffer (20 mM Tris-HCl pH 7.9, 500 mM NaCl and 10 % glycerol) and lysed by sonication at 40 % power, 4 times for 45 seconds (Bandelin Sonopuls). The lysates were centrifuged at 17500 g for 20 min at 4 °C. The supernatant was incubated for 15 min at 4 °C with 5 mL of Ni^2+^-beads (Qiagen Ni-NTA Superflow) equilibrated in lysis buffer. After removal of the flow-through, the beads were washed with 2 column volumes of buffer (20 mM Tris-HCl pH 7.9, 500 mM NaCl and 10 % glycerol) containing 5 mM, 10 mM and 15 mM imidazole, in that order. The elution of the protein of interest was done in a buffer composed of 100 mM Tris-HCl pH 7.9, 500 mM NaCl, 10 % glycerol and 250 mM imidazole in 5 steps of 1 column volume each with 15 min incubation. The eluate was concentrated in Amicon Ultra centrifugal filter units (MWCO: 10 kDa; Cytiva). Following affinity chromatography, the proteins were further purified by size exclusion chromatography using an ÄKTA FPLC system equipped with an S200 10/300 GL or S75 10/300 GL column equilibrated in a buffer composed of 20 mM Tris-HCl pH 7.9 and 150 mM NaCl.

### Preparation of human legumain

Human legumain was expressed, purified and activated based on a protocol previously described (Dall et al., 2022). Briefly, the sequence of human prolegumain was cloned into the pLEXSY-sat2.1 vector and transfected into LEXSY P10 host strain of *Leishmania tarentolae* cells of the LEXSYcon2.1 expression system (Jena Biosciences, Jena, Germany). The resulting construct carried an N-terminal His6-tag and an N-terminal signal sequence for secretion into the supernatant. Cultures were grown in brain heart infusion (BHI) medium (Jena Biosciences) in tissue culture flasks with 5 μg/ml of hemin, 50 units/mL penicillin and 50 units/mL streptomycin (Pen-Strep, Carl Roth) in the presence of nourseothricin (Jena Bioscience) as selection antibiotic. Expression was carried out for 48 hours in the dark at 26 °C and 140 rpm. The supernatant containing the protein of interest was harvested by centrifugation and incubated with Ni^2+^-beads at 4 °C. The washing buffer consisted of 50 mM HEPES pH 7.5, 300 mM NaCl and 5 mM β-mercapto ethanol. The elution buffer was the same as washing buffer with addition of 250 mM imidazole. After elution, the protein was concentrated, and PD-10 columns (GE Healthcare, Uppsala, Sweden) were used for buffer exchange with storage buffer 20 mM HEPES pH 7.0, 50 mM NaCl and 2 mM DTT. The pH-driven auto activation was performed with incubation at room temperature at pH 3.5 after which size exclusion chromatography was performed with buffer composed of 20 mM citric acid pH 4.0, 50 mM NaCl and 2 mM DTT.

### Cleavage assays

Legumain was incubated with SET constructs at a 1:10 molar ratio in a buffer composed of 100 mM citric acid pH 5.5 or 100 mM Hepes pH 7.0 and 150 mM NaCl at 37 °C. Samples were collected after 1.5 hours and subsequently analyzed by SDS-PAGE.

### Cleavage analysis by mass spectrometry

For mass spectrometric analysis, samples were desalted with C18 ZipTips (Merck Millipore), eluted from the tips with 50% acetonitrile in 0.1% formic acid and directly infused into the mass spectrometer (Q-Exactive, Thermo Fisher Scientific) at a flow rate of 1 µl/min. Capillary voltage at the nanospray head was 2kV. Raw data were processed with Protein Deconvolution 2.0 (Thermo Fisher Scientific). Masses were assigned to the protein sequence with the Protein/Peptide Editor module of BioLynx (part of MassLynx V4.1, Waters).

### Analysis of oligomerization state via SEC

To analyze the oligomerization state of uncleaved and cleaved SET variants, 500 µl of sample were loaded on a S200 10/300 GL column equilibrated in a buffer composed of 20 mM Tris pH 7.9 and 150 mM NaCl. For all samples investigated, fractions were collected and analyzed by SDS-PAGE.

### nanoDSF experiments

To access the thermal stability of different SET constructs, we performed differential scanning fluorimetry experiments. The proteins to be analyzed were incubated in assay buffer composed of 20 mM Tris pH 7.9 and 150 mM NaCl supplemented with 100 mM Hepes pH 8.0 or pH 7.0 or BisTris pH 6.5 or citric acid pH 5.5 or pH 4.0 for 5 minutes at a final protein concentration of 1 mg/ml. Suitable capillaries were filled with protein solution (approximately 10 μl) and changes in intrinsic fluorescence intensity were measured at 330 and 350 nm using a Tycho NT.6 instrument (Nanotemper) upon heating the samples from 35 °C to 95 °C within a total measuring time of approx. 3 minutes.

### Protein crystallization and structure solution

First, cleaved SETΔC was prepared by mixing SETΔC and legumain in a 1:100 molar ratio at pH 5.5 for 1 hour at 22 °C. The reaction was stopped by the addition of 1 mM S-methyl-methanethiosulfonate (MMTS). Cleaved SETΔC was subsequently concentrated and subjected to SEC experiments in order to separate cleaved SETΔC from legumain following the procedure described above. Fractions suitable for crystallization were selected following SDS-PAGE and concentrated to a final protein concentration of 70 mg/ml. Initial crystallization screening was performed in a sitting-drop vapour diffusion setup. 0.2 µl of concentrated cleaved SETΔC were mixed with 0.2 µl screen solution (Hampton Index HT or Hampton Crystal Screen I + II) and equilibrated with 70 µl reservoir solution in 96-well MRCII plates (SwissCI, Zug, Switzerland) at 20 °C. Crystallization reactions were set up using a Mosquito Xtal3 crystallization robot (sptlabtech, Melbourn, England) and analysed using a RockImager 1000 imaging system (Formulatrix, Dubai, United Arab Emirates). After one week, crystals were observed in a condition composed of 26 % PEG 400, 0.2 M calcium chloride dihydrate and 0.1 M Hepes pH 6.75. Crystals were harvested and flash-frozen in liquid nitrogen. A native dataset was collected at 100 K on beamline ID30A3 (ESRF, Grenoble) equipped with an Eiger X 4M detector to a resolution of 2.47 Å. 3600 images were collected at a wavelength of 0.967 Å at 0.1 ° oscillation range. Xray diffraction data was processed utilizing XDS (34). An initial model was obtained by molecular replacement using AutoMR from the Phenix (Python-based Hierarchical Environment for Integrated Xtallography) program suite utilizing coordinates of SET (pdb 2e50). Iterative cycles of rebuilding in COOT (35) followed by refinement in phenix.refine (36) and REFMAC (37) were carried out. Coordinates and structure factors were deposited with the PDB under entry code 9I15. Pymol (38) was used to create figures illustrating structures.

### Histone binding assay

A mixture of histones H1, H2A, H2B, H3 and H4, isolated from calf thymus was purchased (Merck, Darmstadt, Germany). The histone mix was diluted in PBS to a concentration of 10.44 mg/ml. The histone pull-down assay was performed using SET-FL, SETΔC, and SET-FL and SETΔC after cleavage by legumain, following a protocol described by Muto S. et al. (28). All SET constructs harbored a C-terminal His_6_-tag, which was exploited for Ni^2+^-affinity purification. Specifically, 0.5 mg/ml of the respective SET construct were mixed with 1 mg/ml histone mix in assay buffer containing 20 mM Hepes pH 8.0, 300 mM KCl, 20 % glycerol, 0.05 % Tween 20 and 5 mM imidazole and incubated for 50 min at 4 °C. Subsequently, the reaction mix was incubated with Ni^2+^-beads preequilibrated in assay buffer for 30 min at 4 °C. The fraction containing unbound protein was isolated upon centrifugation at 2000 g for 2 min at 4 °C. Beads were washed 3 times with assay buffer followed by one wash with assay buffer supplemented with 10 mM and 15 mM imidazole respectively. Bound protein complexes were eluted using assay buffer supplemented with 250 mM imidazole. The elution step was repeated 4 times. Control reactions contained histones only or SET proteins only respectively. Fractions were analyzed via SDS-PAGE.

### Microscale thermophoresis (MST) experiments

To test the interaction of intact and cleaved SET-FL and SETΔC proteins with histone H1, microscale thermophoresis (MST) experiments were performed. 100 µl of bovine histone 1 or human histone 1 (Merck, Darmstadt, Germany) at a concentration of 20 µM were buffer exchanged into labeling buffer composed of 100 mM Hepes pH 8.0 and 50 mM KCl. Subsequently histones were labeled with NT-647-NHS dye (Nanotemper) at a molar ratio of 3:1 (protein:dye) in labeling buffer for 30 min. at 20 °C in the dark. Unreacted, excess dye was removed using a NAP-5 column (GE Healthcare) preequilibrated in PBS buffer. Fractions of 100 µl were collected and DOL (degree of labeling) and protein concentration were determined. To determine the binding affinity of the different SET variants for bovine histone 1, a 1:2 dilution series was prepared in assay buffer composed of 20 mM Hepes pH 8.0, 150 mM KCl and 0.05 % Tween-20 ranging from 32 µM to 0.8 nM. Labeled bovine histone 1 was diluted in assay buffer to reach a final concentration of 14.4 nM. 10 µl of the respective SET dilution were mixed with 10 µl labeled histone H1 and incubated for 15 min at 20 °C in the dark. To remove possible aggregates, the complexes were centrifuged at 13.000 g and 20 °C for 10 min. Immediately after centrifugation, the reactions were transferred into standard coated capillaries and MST traces were measured at 25 °C, 40% LED and medium MST power in an NT.115 RED instrument. Similar MST experiments were carried out using labeled human histone H1 at a concentration of 11 nM. Ligand binding was investigated by analyzing changes in MST traces. Data were recorded with the MO.Control 1.5.1 (NanoTemper) software and further analyzed using MO-Affinity Analysis 2.2.6 (NanoTemper). MST traces were processed using an MST on-time of 5 sec and fitted using Kd (assuming 1:1 binding) fit models. Kd values were determined from three independent binding experiments. To confirm that the shift in MST and/or fluorescence signal was due to complex formation rather than unspecific aggregation, samples were incubated with 10% SDS and 10 mM DTT and heated to 95 °C for 5 min before measuring MST traces.

### PP2A assay

HEK293T cells were lysed by sonication in a buffer composed of 20 mM imidazole pH 7.0, 2 mM EDTA, 2 mM EGTA and 1 % protease inhibitor cocktail (P8340, Merck, Darmstadt, Germany). The soluble fraction was obtained by centrifugation at 13000 rpm for 10 min at 4 °C. 100 µg of protein were mixed with 4 µg of anti-PP2A antibody C subunit clone 1D6 (Merck, Darmstadt, Germany) and 40 µl of rec-Protein G-Sepharose® 4B Conjugate (Invitrogen, Camarillo, USA). The volume was filled up to 500 µl using assay buffer composed of 50 mM Tris pH 7.0 and 100 µM CaCl_2_. Reactions were incubated for 1 hour at 4 °C with constant rocking. Beads were harvested by centrifugation at 2000 rpm, 20 °C for 2 min and subsequently washed with TBS for 5 min at 20 °C under constant rocking. Subsequently, 3 µM of the respective SET construct was added and incubated for 10 min at 20 °C. In control experiments assay buffer was added instead of the SET proteins. Then, 750 µM of K-R-pT-I-R-R phosphopeptide (Merck, Darmstadt, Germany) were added and again incubated for 10 minutes at 30 °C shaking at 750 rpm. The reaction volume was adjusted to 80 µl using assay buffer and 20 µl malachite green phosphate detection solution (Merck, Darmstadt, Germany) was added. Color was developed for 30 min. at 20 °C and absorbance was measured at 620 nm using a CLARIOstar Plus microplate reader (BMG Labtech, Ortenberg, Germany).

### Alphafold modelling

A model of SET-FL was obtained using a local installation of Alphafold 2. The sequence used for modeling comprised residues Met1-Asp277 according to Uniprot accession code Q01105-2. The PyMOL Molecular Graphics System (Schrödinger, LLC) was used to illustrate protein models.

## Accession numbers

The crystal structure of cleaved SETΔC was deposited with the wwPDB under the entry code 9I15.

## Acknowledgements

The authors wish to thank Martina Wiesbauer for technical assistance and the Austrian Science Fund (FWF, project number Y1469 to E.D.) for funding.

