## Supplementary Information for "Conformational and Functional Regulation of SET by Legumain Cleavage"

### Supplementary Figure 1

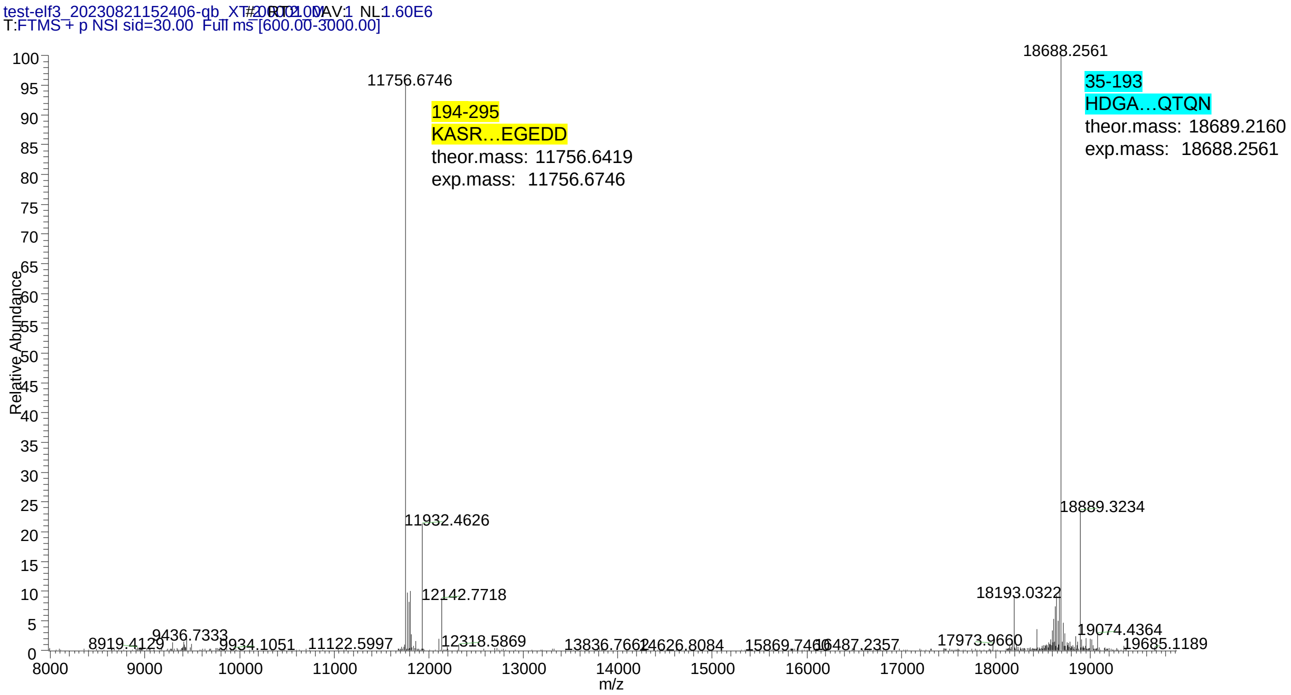


**Sequence of Full-length SET**

**MGHHHHHHSGTENLYFQGMSAPAAKVSKKELNSNHDGADETSEKEQQEAIEHIDEVQNEIDRLNEQASEEILKV**
**EQKYNKLRQPFFQKRSELIAKIPNFWVTTFVNHPQVSALLGEEDEEALHYLTRVEVTEFEDIKSGYRIDFYFDENPYF**
**ENKVLSKEFHLNESGDPSSKSTEIKWKSGKDLTKRSSQTQNKASRKRQHEEPESFFTWFTDHSDAGADELGEVIKD**
**DIWPNPLQYYLVPDMDDEEGEGEEDDDDDEEEEGLEDIDEEGDEDEGEEDEDDDEGEEGEEDEGEDD**

**Supplementary Figure 1. SET-FL is processed by legumain at two major sites.** Intact mass of SET-FL following cleavage by legumain revealed two major cleavage products indicated in cyan and yellow respectively. The cleavage reaction was set up at pH 5.5. The major cleavage sites are Asn16 on the N-terminal region and Asn175 on the earmuff domain.

### Supplementary Figure 2

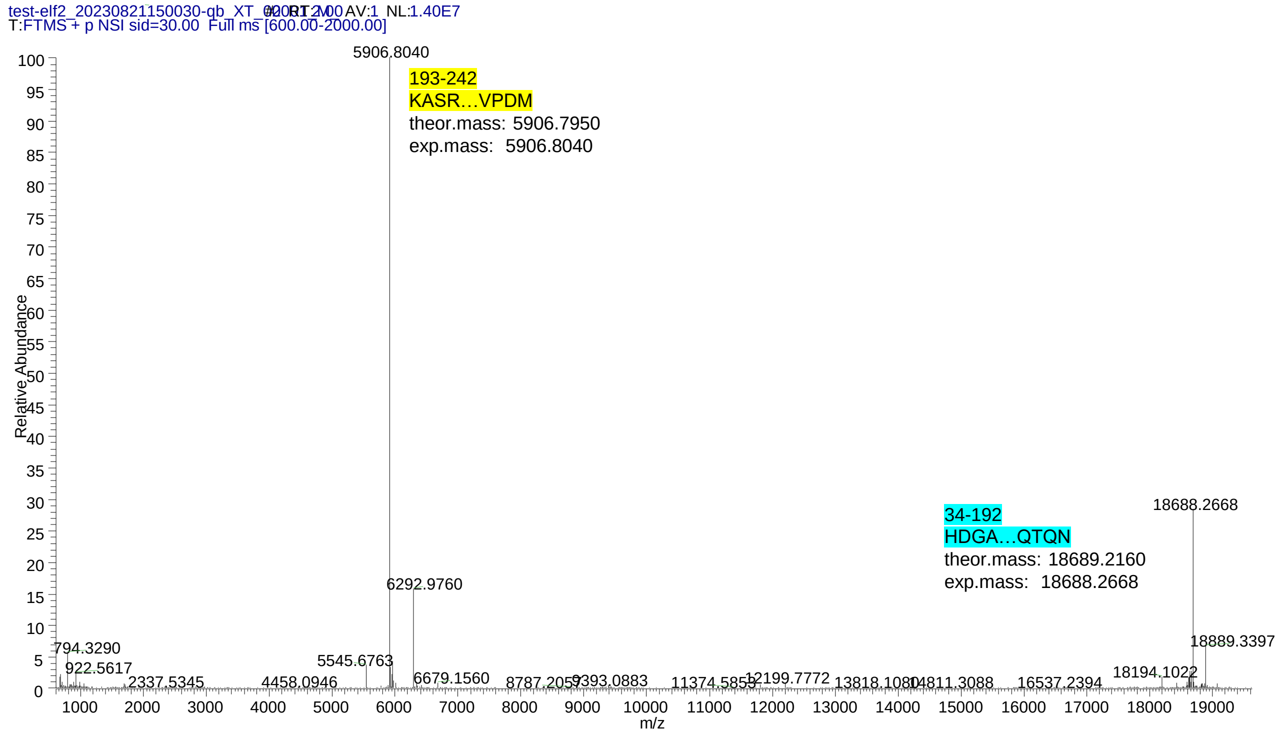


**Sequence of SETΔC**

MGHHHHHHSGTENLYFQGSAPAAKVSKKELNSNHDGADETSEKEQQEAIEHIDEVQNEIDRLNEQ
ASEEILKVEQKYNKLRQPFFQKRSELIAKIPNFWVTTFVNHPQVSALLGEEDEEALHYLTRVEVTEFEDI
KSGYRIDFYFDENPYFENKVLSKEFHLNESGDPSSKSTEIKWKSGKDLTKRSSQTQNKASRKRQHEE
PESFFTWFTDHSDAGADELGEVIKDDIWPNPLQYYLVPDM

**Supplementary Figure 2. SETΔC is processed by legumain at two major sites.** Mass spectrum of a direct infusion mass spectrometry experiments revealed two major cleavage products indicated in cyan and yellow respectively. The cleavage reaction was set up at pH 5.5. The major cleavage sites are Asn15 on the N-terminal region and Asn175 on the earmuff domain.

### Supplementary Figure 3

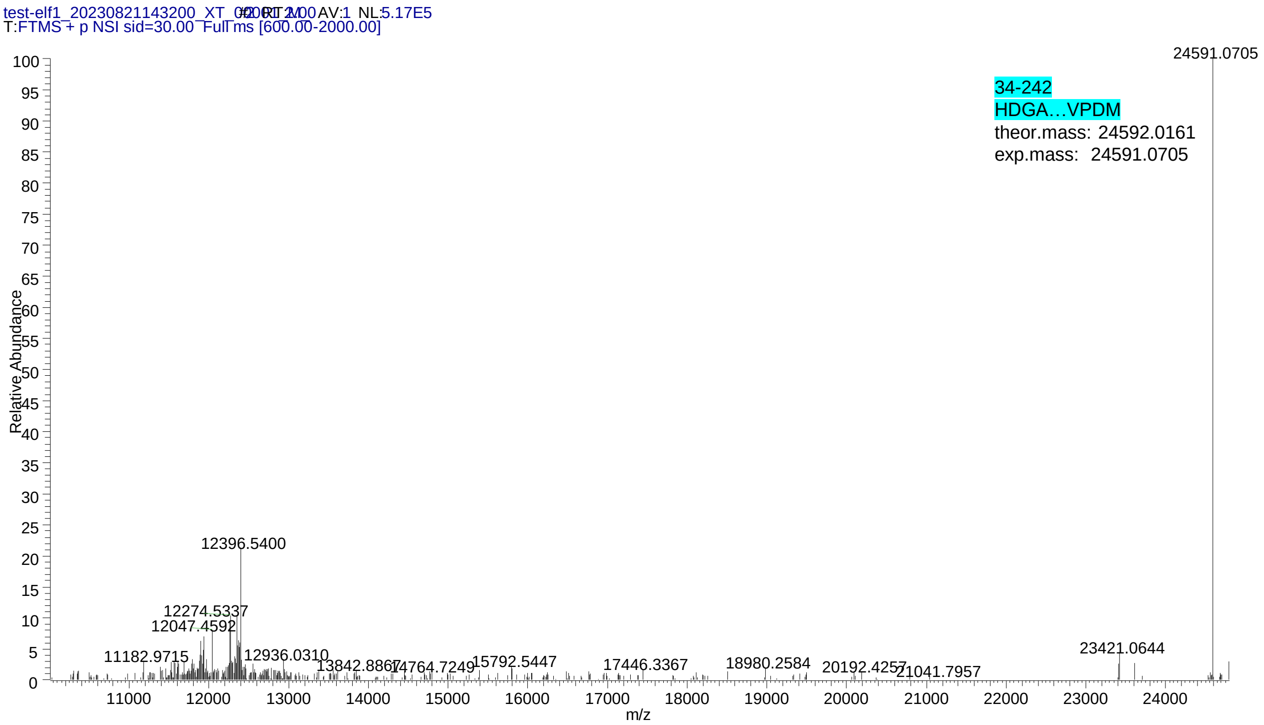


**Sequence of SETΔC-N175Q**

**MGHHHHHHSGTENLYFQGSAPAAKVSKKELNSNHDGADETSEKEQQEAIEHIDEVQNEIDRLNEQ ASEEILKVEQKYNKLRQPFFQKRSELIAKIPNFWVTTFVNHPQVSALLGEEDEEALHYLTRVEVTEFEDI KSGYRIDFYFDENPYFENKVLSKEFHLNESGDPSSKSTEIKWKSGKDLTKRSSQTQQKASRKRQHEE PESFFTWFTDHSDAGADELGEVIKDDIWPNPLQYYLVPDM**

**Supplementary Figure 3. SETΔC-N175Q is not processed by legumain after residue 175.** Mass spectrum of a direct infusion mass spectrometry experiments revealed one major cleavage site, Asn16 on the N-terminal region. The cleavage reaction was set up at pH 5.5.

### Supplementary Figure 4

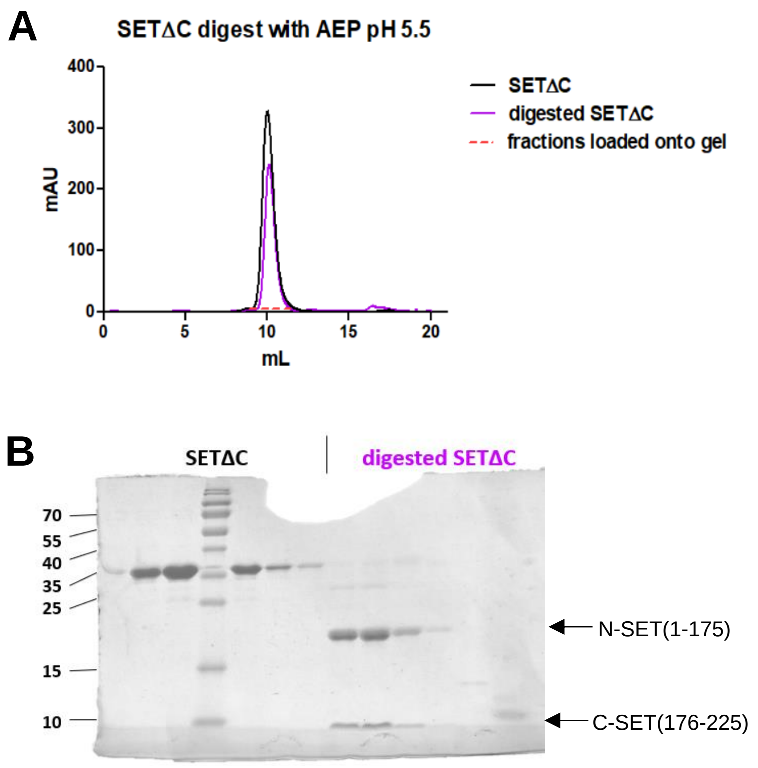


**Supplementary Figure 4. N- and C-terminal cleavage products of SETΔC do not dissociate after cleavage by legumain. A)** Size exclusion chromatography experiment of intact and cleaved SETΔC at pH 5.5. **B)** Peak fractions of the experiment shown in A) were analyzed by SDS-PAGE.

**Supplementary Figure 5**

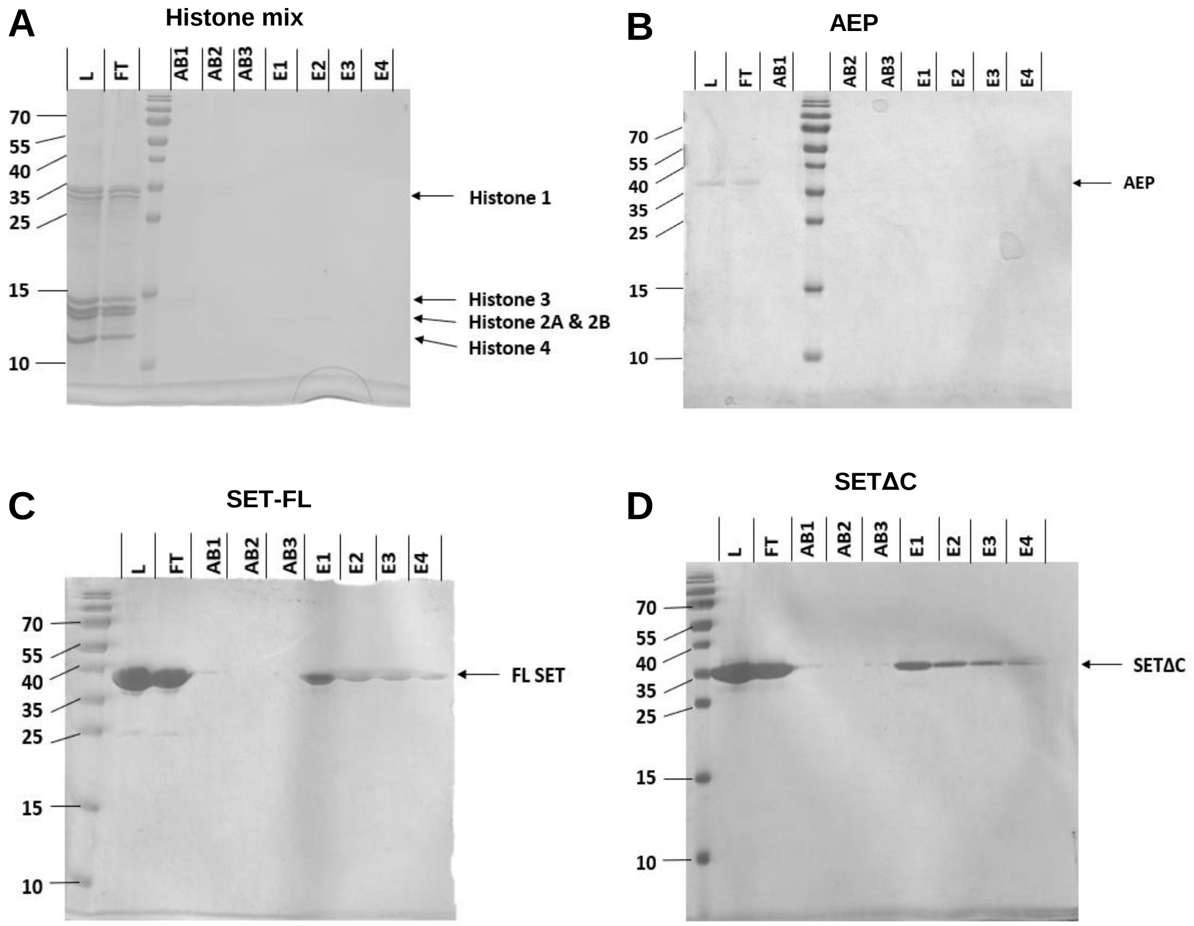


**Supplementary Figure 5. Control experiments revealing that histones and AEP do not bind to $Ni^{2+}$-beads, and SET-FL and SETΔC do bind.** A mix of bovine histones (**A**), AEP (legumain) (**B**), SET-FL (**C**) or SETΔC (**D**) were subjected to a $Ni^{2+}$-purification and fractions were analyzed by SDS-PAGE. L: load, FT: flow through, AB1-3: wash fractions 1-3, E1-4: elution fractions 1-4.

### Supplementary Figure 6

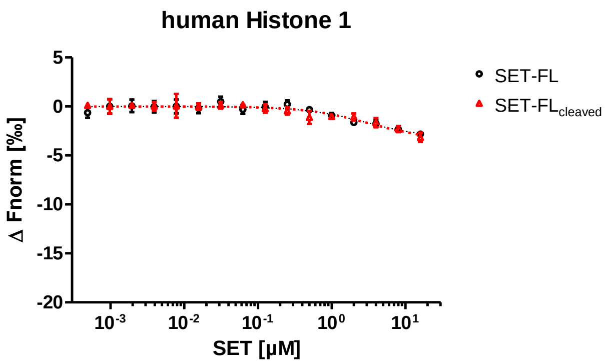


**Supplementary Figure 6. Intact and cleaved SET-FL bind similarly to human histone H1.** Microscale thermophoresis (MST) experiments testing the interaction of indicated SET variants to human histone 1.
